# FERMO: a Dashboard for Automated Prioritization of Molecular Features from Mass Spectral Data

**DOI:** 10.1101/2022.12.21.521422

**Authors:** Mitja M. Zdouc, Hannah E. Augustijn, Nataliia V. Machushynets, Lina M. Bayona, Sylvia Soldatou, Niek F. de Jonge, Sarah Casu, Marcel Jaspars, Gilles P. van Wezel, Marnix H. Medema, Justin J. J. van der Hooft

## Abstract

Small molecules shape phenotypic variation by modulating biological processes, yet linking specific molecules to observed traits remains challenging due to the complexity of biological samples. Liquid chromatography-tandem mass spectrometry routinely detects hundreds of molecules per sample, and computational tools aid in the selection of the biologically relevant subset by organizing, annotating, and integrating orthogonal data. Existing tools typically focus on facilitating data-driven exploration to support manual interpretation, rather than more objective, data-driven prioritization and hypothesis-generation. Here, we introduce FERMO, a free online dashboard interface for prioritization of molecular features and samples associated with phenotypes of interest. FERMO’s modular framework automates data processing, annotation, and integration of standardized phenotypic and other metadata. FERMO supports both exploratory and targeted analysis through efficient interactive visualization, reproducible prioritization, and data filtering. We demonstrate FERMO’s utility in benchmarking studies prioritizing bioactive compounds from complex biological matrices. FERMO is freely available at https://fermo.bioinformatics.nl/.

## Introduction

Small molecules are an integral component of the environment. By selectively interacting with biological targets and systems, they play a crucial role in driving phenotypic variance^1^. Therefore, their investigation is central to fields such as natural product drug discovery, exposomics, or microbiome research^2^. Furthermore, such specialized metabolites often give an evolutionary advantage to the producing organism^3^, and have yielded a wide range of drugs, pesticides, and industrial applications^4^.

Liquid chromatography - mass spectrometry (LC-MS) is a widely used analytical method in untargeted metabolomics, offering unparalleled sensitivity and high throughput^5^. It detects so-called mass or molecular/metabolite features (from here, features), which are ionized molecules characterized by their specific mass-to-charge ratio (*m/z*) and column retention time (RT). A single molecule can be associated with multiple ion species, resulting in several features representing the same molecule, whereas non-ionized molecules are not detected by mass spectrometry^2^. Additionally, ionized molecules can be fragmented through tandem mass spectrometry (MS/MS), generating fragment ions that are determined by the chemical structure of their parent molecule, thus providing a “molecule fingerprint” for identification^6^. Untargeted LC-MS/MS metabolomics allows for a comprehensive capture of a sample’s chemical composition, routinely detecting hundreds of features per sample. However, the complexity of these datasets makes manual analysis infeasible in most cases^7^. To address this, metabolomics software tools have been developed to process LC-MS/MS data and streamline the metabolomics analysis workflow. These tools generally include functionalities such as feature organization, annotation, and data visualization, both within graphical user interfaces (GUIs) and command line interfaces (CLIs)^8–15^. Most tools are primarily designed for untargeted, exploratory data analysis and not optimized for hypothesis-driven prioritization of samples and metabolite features therein^16^. Such more targeted analysis typically requires integration of orthogonal data, like phenotypic data, to guide selection of molecules relevant to the research question. A few applications have been designed specifically for hypothesis-driven prioritization by automatically integrating orthogonal data^14,17,18^. With regard to phenotypic data, previous efforts resulted in Bioactivity-Based Molecular Networking^19^ and Bioactive Natural Products Prioritization^20^, which use custom scripts to overlay spectral networks generated through the GNPS web application^11^ with biological activity and bibliographic metadata. More recently, several CLI tools have been developed to correlate feature abundance with biological readouts, but they lack GUIs^21,22^. Another tool is NP Analyst, an online GUI prioritization tool for correlating LC-MS/MS metabolomics with high-throughput phenotype screening data by calculating Activity and Cluster Scores^23^. While this and other tools effectively integrate metabolomics with orthogonal data, their scope is typically limited to specific tasks or scenarios within the overall metabolomics prioritization workflow. Hence, a tool that offers systematic, flexible, and reproducible prioritization of relevant samples and metabolite features in untargeted mass spectrometry profiles linked to a GUI is lacking.

To address this gap, we developed FERMO, a free open-source web application that provides a generalized, comprehensive workflow for hypothesis-driven prioritization. FERMO automatically integrates metabolomics with phenotypic, genomic, and sample metadata, and performs feature organization and annotation for both positive and negative ion modes. By automating the calculation and aggregation of key molecular information into feature and sample scores, FERMO streamlines the prioritization process. Results are presented through a GUI dashboard, with a customizable filter panel enabling users to focus on the most relevant features and samples and facilitates data-driven decision-making. We demonstrate the capability of the tool by benchmarking FERMO on two case studies, re-analyzing previously reported, experimentally validated datasets. FERMO is designed for workflow reproducibility, employing standardized input and output formats and structured input parameter management. Thus, it enhances hypothesis-driven metabolomics prioritization across various fields, including natural product research, environmental science, and microbiome and agricultural research.

## Results

### Hypothesis-driven prioritization using the FERMO dashboard

Metabolomics data analysis routinely yields thousands of features, obfuscating the identification of biologically relevant signals. Therefore, efficient integration of orthogonal data capturing both causes and consequences of phenotypic variation is essential for hypothesis-driven prioritization. To this end, FERMO was developed as a framework for automated data integration and prioritization of candidate molecules from complex biological matrices. An initial beta version of FERMO was released in 2022^24^, shaped by extensive scientific exchanges with the metabolomics community and insights from earlier prototypes and use cases^25–28^ in order to facilitate early access to the technology and benefit from further community feedback. We then made a further refined FERMO dashboard (version 1.0) available as a web application in 2024, incorporating input from both academic and industry participants collected during a structured beta-testing pilot study (Table S1A).

FERMO revolves around an interactive, dynamically updated dashboard-style GUI, which provides a convenient cross-sample overview of feature information through interactive and freely arrangeable panels, each focusing on different data representations (Fig.1). Following the “overview first - details on demand” principle^16^, the sample overview (Fig. 1a) allows users to select samples for chromatogram visualization (Fig. 1b) based on descriptive attributes and calculated scores. Features are represented as pseudo-chromatograms, approximations of extracted ion chromatograms (EICs) constructed from discrete data points (for further details see Supplementary Methods and Fig. S1). Selection of pseudo-chromatograms toggles detailed information and annotations on features within and across samples (Fig. 1c) and spectral similarity networking information as graph- and chromatogram views (Fig. 1d). The filter panel (Fig. 1e) allows for cross-sample feature prioritization with the help of eleven different filter settings, including phenotype information, presence across groups, and annotation status. This allows for rapid identification of features and samples of interest. All results can be downloaded, including the annotated peak table and full spectral networks, formatted for import into Cytoscape^29^. An extensive documentation including full description of expected input and generated output files as well as explanatory videos for most steps can be found in the Documentation (Table S1B).

**Fig. 1.**
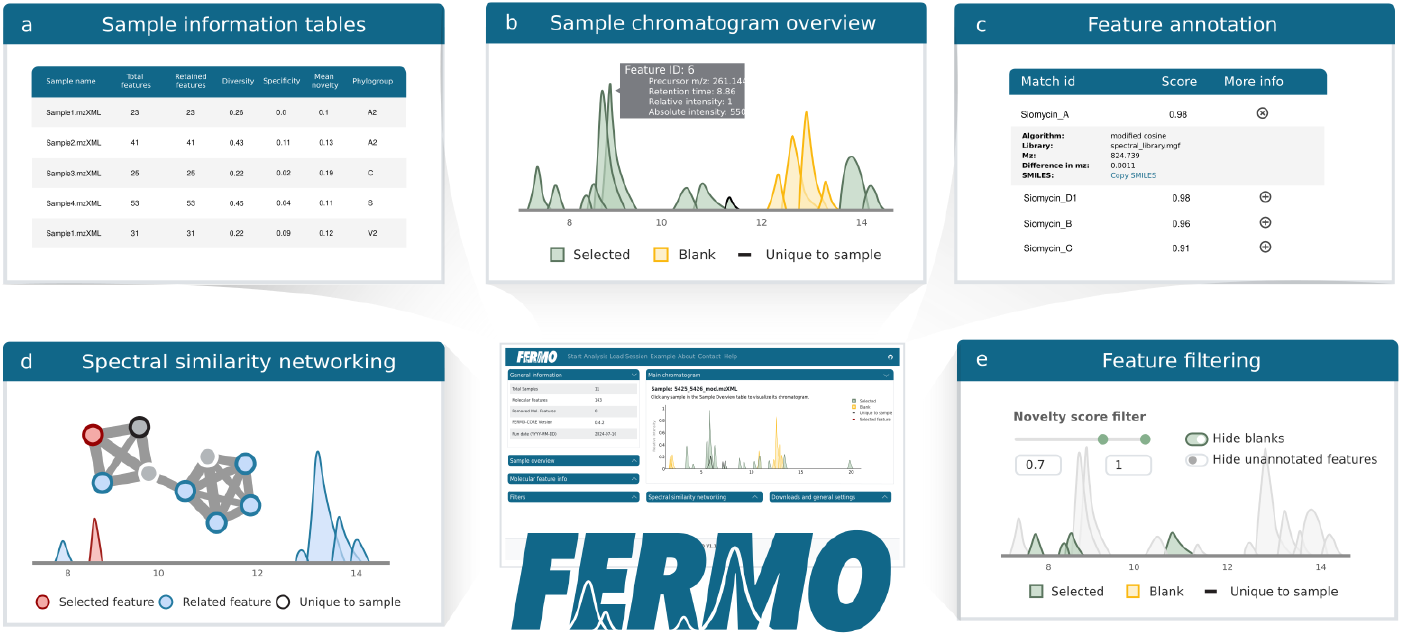
FERMO dashboard overview. Functionality of the FERMO dashboard is depicted in the individual panels. **a)** Sample information tables displaying the total number of detected features, retained features at current filter settings, calculated scores, and sample metadata; **b)** sample chromatogram visualizations, with approximations of EICs as pseudo-chromatograms constructed from discrete data points such as RT and feature-width-at-half-maximum (FWHM). Features retained at the current filter settings shown as green peaks, sample blank-associated features in yellow, and sample-unique features with black outlines; **c)** feature information visualized in drop-down panels, including ion identity, fragment and neutral loss annotations, library matching, group details, and cross-sample intensity and area; **d)** cross-sample spectral similarity networking representation, with the pseudo-chromatogram showing features present in the currently inspected sample; **e)** Feature prioritization using a combination of up to eleven filtering options, including phenotype and novelty scores, annotations, precursor *m/z* range, and presence in specific sample groups.

### Modular orthogonal data integration with FERMO

FERMO is designed as a modular and flexible application, with a focus on orthogonal data integration. Besides a mandatory peak table, it accepts any combination of MS/MS spectrum, phenotype information, sample grouping, and spectral library files (Fig. 2a). Additionally, it accepts results from MS2Query^30^ and genomic analysis data retrieved from the biosynthetic gene cluster (BGC) prediction tool antiSMASH^31^. While a default selection of activated modules and parameters settings are provided for convenience, users may freely select modules and adjust parameters suitable for their data and research question. Of note, providing a more comprehensive set of input files enhances the analysis quality, improves annotation depth, and facilitates more effective prioritization. Once analysis is initiated, FERMO’s processing backend (fermo_core) performs data processing in the form of an automated analysis pipeline, including input processing, data organization, annotation, and score calculation (Fig. 2b). The prioritization scores are key to rapid, hypothesis-driven prioritization using the dashboard interface, and integrate results from multiple analysis modules. Feature-associated scores include the Phenotype Score, which reflects the highest correlation between a feature and integrated phenotype data, and the Novelty Score, which estimates the putative novelty of a molecule relative to reference spectral libraries. Sample-associated scores include the Mean Novelty Score, representing the average novelty of all features of a given sample, the Diversity Score, which quantifies the chemical diversity of a sample in comparison to others in the dataset, and the Specificity Score, which measures the fraction of chemical diversity unique to a single sample (Supplementary Methods). All analysis results, input file names, and parameter settings are written to FERMO Session output files. These Javascript Object Notation (JSON) files serve as primary output of FERMO, from which the dashboard visualization is directly loaded (Fig. 2c). A preliminary benchmarking showed that FERMO can process and load datasets of several thousand features in a reasonable time (Table S2). FERMO Session files follow a compact, structured data schema and can be downloaded for long-term storage and conveniently exchanged between collaborators. Of note, FERMO run parameters can be directly loaded from session files, facilitating reproducible analysis. Therefore, we recommend users of FERMO to always deposit Session files alongside reported results, similar to the Batch files generated by mzmine for reproducible data analysis^32^.

**Fig. 2.**
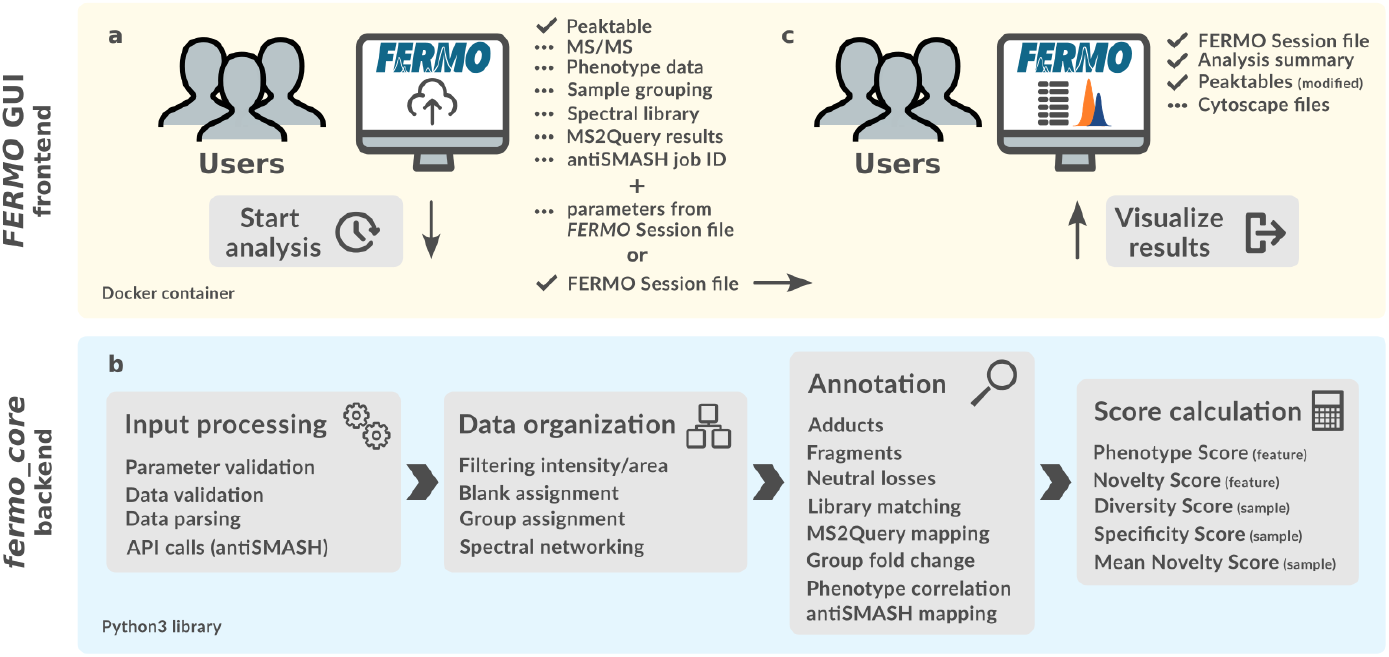
Schematic overview of data analysis steps conducted by FERMO. Functionality of the tool can be separated in the FERMO frontend and the fermo_core backend. **a)** Users can submit jobs using the FERMO GUI app by uploading a peak table (mandatory) and any combination of additional files (optional). While default parameters are provided, these can be freely changed, or loaded from a previously generated FERMO Session file. Previous FERMO jobs can be uploaded as well, directly leading to c). **b)** Data processing is performed by the fermo_core backend processing pipeline, with module execution dependent on user input and parameter settings. **c)** After completion, results and parameter settings are written in a FERMO Session file, from which the dashboard visualization is loaded. All generated files can be downloaded, shared, and used for follow-up analyses.

### Benchmarking FERMO against established phenotype-prioritization workflows

To evaluate FERMO’s performance in hypothesis-driven prioritization, we benchmarked its phenotype scoring module against established workflows. While FERMO uniquely integrates phenotype data with MS/MS annotation, sample grouping, and genomics information within a modular, dashboard-driven framework (Table S3), individual modules can be directly compared to existing tools. We selected a publicly available dataset derived from the plant *Euphorbia dendroides*, consisting of crude extracts and chromatographic fractions with associated antiviral activity against chikungunya virus (MassIVE ID: MSV000080502). This dataset, originally analyzed with the Bioactivity-Based Molecular Networking (BioMN) approach^19^, has since become a reference point for bioactivity-guided prioritization. We compared FERMO’s results with BioMN and the recently published NP3 MS Workflow^22^. NP Analyst^23^ was excluded from the analysis due to its requirement for multiple phenotypic assays, which the dataset does not meet.

Re-analysis of the dataset using mzmine3 yielded 912 features detected in two or more samples. After removing low-abundance features (≤1% of relative area in any sample) using FERMO’s built-in filtering tools, 737 features remained - comparable to the 587 features reported by the original BioMN analysis. Using FERMO’s phenotype scoring module (Pearson correlation with r ≥ 0.85 and false discovery rate (FDR)-adjusted *p* ≤ 0.05), we successfully recovered the same three experimentally validated features as BioMN (Features **20–22**; *m/z* 591.324 at 27.05 min; 589.311 at 25.4 min; 563.296 at 21.6 min; see Fig. 3). A fourth compound (**23**), which had been isolated but not prioritized by BioMN, was also not prioritized by FERMO. In contrast, the NP3 MS Workflow failed to prioritize any of the experimentally validated features, reporting low “bioactivity correlation coefficient scores” (NP3 MS *r*, Fig. 3). As the NP3 MS Workflow does not report FDR-corrected *p*-values, it was excluded from further comparison.

**Fig. 3.**
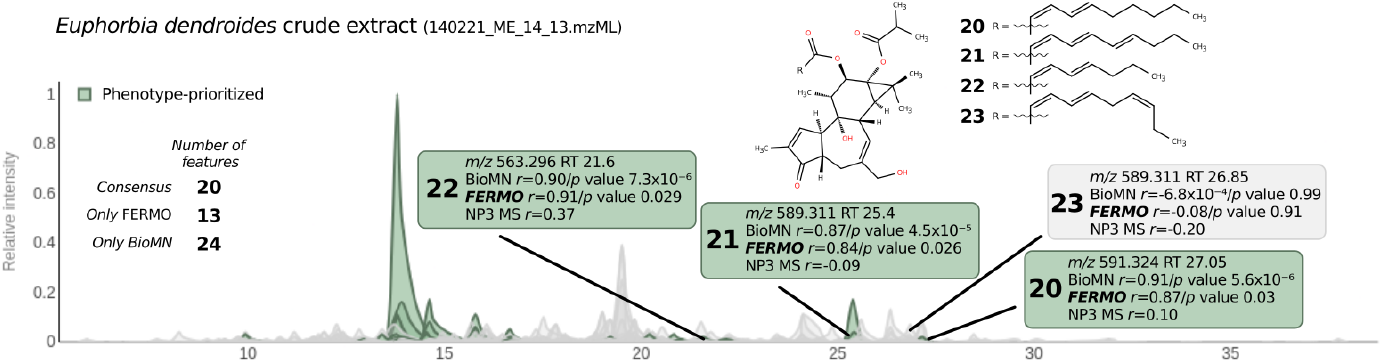
Benchmarking of phenotype-prioritization workflows. FERMO is benchmarked against Bioactivity-Based Molecular Networking^19^ and the NP3 MS Workflow^22^, visualized using FERMO’s pseudo-chromatogram view. Features prioritized by FERMO based on phenotype score are highlighted in green; features excluded by the phenotype score filter are shown in grey. Both FERMO and BioMN prioritized a comparable set of features, including compounds **20–22**, which were experimentally validated in the original BioMN study. Feature **23**, which was isolated but not prioritized in the BioMN study, was also not prioritized by FERMO. In contrast, the NP3 MS Workflow reported low correlation scores and no corrected *p-*values for the discussed features and did not identify any as phenotype-associated.

Quantitatively, FERMO and BioMN showed comparable performance, with 20 overlapping prioritized features, 13 unique to FERMO, and 24 unique to BioMN (Table S1C). However, when inspecting statistical significance, FERMO consistently yielded higher (less significant) FDR-adjusted *p*-values. To investigate this, we compared the correlation algorithms and identified a key methodological difference: while both tools use Pearson correlation, FERMO calculates correlations referencing only samples where a feature was detected (non-zero, non-null intensity), whereas BioMN includes all samples, assigning zero intensity where a feature was not detected. This difference significantly impacts the effective sample size and *p*-value calculation (see Table S1D for a detailed comparison via Jupyter notebook).

FERMO also provides additional flexibility and transparency: users can select from multiple FDR correction methods, adapt the algorithm to quantitative, semi-quantitative, or binary phenotype data, and retain all settings and results in a reproducible session file (Table S1E). This benchmark demonstrates that FERMO not only reproduces key findings of prior studies, but also offers enhanced functionality, user control, and workflow transparency. This enables robust, reproducible, and scalable phenotype-driven prioritization within a single, modular framework.

### FERMO effectively prioritizes bioactive actinomycin in an OSMAC study

To further demonstrate FERMO’s utility in a real-world context, we revisited a previously published study that employed the OSMAC (One Strain, Many Compounds) approach^33^ to elicit specialized metabolite production in *Streptomyces* sp. MBT27 under varying carbon sources^34^. The original study identified actinomycin D as the major bioactive compound through a manual combination of statistical analysis, manual dereplication, and targeted experimental validation. Our aim was to assess whether FERMO could independently prioritize the same bioactive molecule using its automated, hypothesis-driven pipeline.

As the original study did not include a high-resolution LC-MS/MS dataset, we reanalyzed the extracts using high-resolution instrumentation. FERMO was then applied to integrate the resulting metabolomics data with quantitative bioactivity data, as well as with genomic information obtained via antiSMASH analysis of the *Streptomyces* sp. MBT27 genome (GenBank accession NZ_CP070733.1) and annotation results obtained from MS2Query analysis. From a dataset comprising 1,136 features across 36 samples, FERMO prioritized eight features that strongly correlated with the bioactivity profile of the crude extracts (Fig. 4). Six of these were annotated by FERMO as actinomycin D, based on combined results from MS2Query and the antiSMASH KnownClusterBlast module, which linked them to the MIBiG-deposited actinomycin D biosynthetic gene cluster (BGC0000296)^35^. Four of these features, *m/z* 1255.6353, 1277.617, 639.3141, and 628.3233, were dereplicated as the [M+H]^+^, [M+Na]^+^, [M+Na+H]^2+^, and [M+2H]^2+^ ion adducts of actinomycin D, respectively. The remaining two features, *m/z* 1293.612 ([M+Na]^+^) and 636.3205 ([M+2H]^2+^), were dereplicated as the hydroxylated congener actinomycin X_0β_. These findings align exactly with the results reported in the original publication^34^. In addition to reproducing previously validated findings, FERMO highlighted two additional minor features (*m/z* 662.2726 and *m/z* 656.2845), that remain to be structurally characterized, suggesting the potential for novel discovery. Notably, FERMO achieved these results while prioritizing only 0.7% of the dataset (8 out of 1,136 features), underscoring its ability to integrate orthogonal data types, perform accurate annotation and dereplication, and streamline the analysis without requiring pre-fractionated samples. This significantly reduces experimental cost and increases throughput, offering a powerful alternative to traditional data processing workflows.

**Fig. 4.**
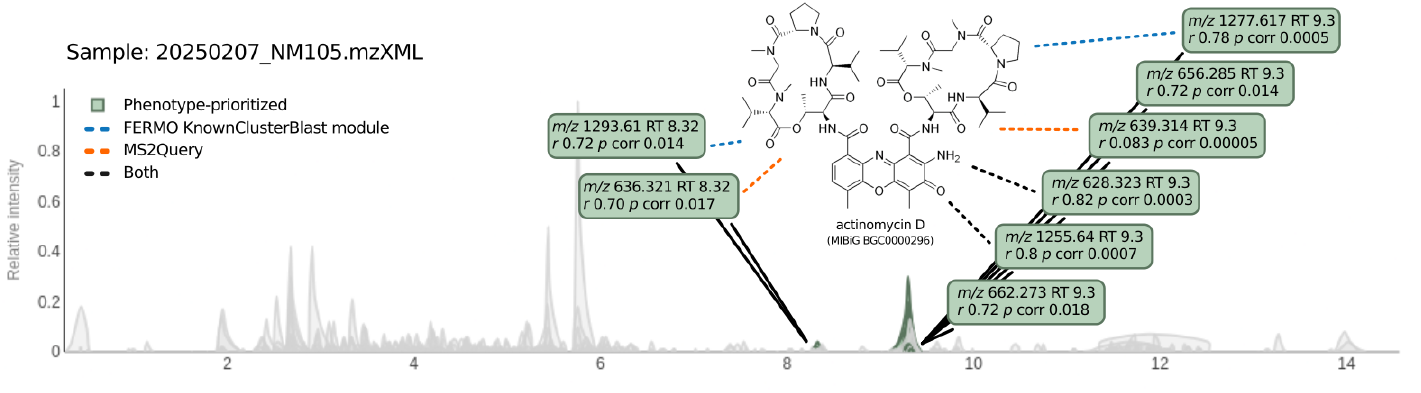
Results of phenotype-based prioritization of samples of *Streptomyces* sp. MBT27. Prioritized features are highlighted in green and annotated with identity annotations, Pearson correlation coefficients, and Bonferroni-corrected *p*-values. Dashed lines indicate features annotated as actinomycin D-related compounds using MS2Query and/or dereplication via the antiSMASH KnownClusterBlast module, referencing the MIBiG database. Ions with *m/z* 1255.6353, 1277.617, 639.3141, and 628.3233 were identified as various adducts of actinomycin D, while annotations for features with *m/z* 1293.612 and 636.3205 were refined as actinomycin X_0*β*_, a hydroxylated congener of actinomycin D. Two additional minor features with *m/z* 662.2726 and *m/z* 656.2845 remain uncharacterized.

## Discussion

FERMO streamlines hypothesis-driven feature prioritization, allowing researchers to allocate resources more efficiently by focusing on compounds most relevant to the respective research question. By integrating orthogonal data and combining both intrinsic properties and derived attributes, FERMO offers a rational, broadly applicable framework for prioritization. Its effectiveness was demonstrated through benchmarking against BioMN and the NP3 MS workflow, as well as in a real-world case study, where FERMO successfully identified bioactivity-associated features previously validated through experimental work. While FERMO does not replace the need for downstream validation, it substantially simplifies and accelerates the interpretation of complex metabolomics datasets.

At its core, FERMO’s dashboard serves as a graphical interface for rapid data exploration, allowing users to inspect and filter features based on prioritization scores that summarize results from individual analysis modules. While several other applications have introduced prioritization scores^18,19,23,25,36^ or dashboard views for metabolomics data analysis^14,37^, FERMO is unique in its comprehensive, systematic modular framework that unifies these elements into a single, fully integrated pipeline for hypothesis-driven prioritization. Among existing tools, NP Analyst^23^ is the most comparable, focusing on integrating LC-MS data with multiple phenotypic assays. However, FERMO distinguishes itself by offering a broader, more flexible solution that extends beyond NPAnalyst’s specific use case in natural product drug discovery. We recognize the parallel development of alternative approaches to represent chemical novelty^38^, diversity^39^, and chemotaxonomic information^36,40^. FERMO is designed as an expandable and modular system, envisioned as the central node within an ecosystem of interoperable tools, similar to the role of the antiSMASH platform in genome annotation^31^. To this end, we encourage developers of complementary prioritization tools to integrate their applications into FERMO as modules or provide web APIs for seamless data exchange. At the same time, we pledge to facilitate long-term maintenance of FERMO and its sustained development. While FERMO is currently supported by institutional infrastructure, its containerized design allows deployment on any other platform, as well as local installation under a permissive open-source license. The development and maintenance of FERMO are overseen by a governing body within the FERMO Metabolomics GitHub Organization (https://github.com/fermo-metabolomics), following O3 and FAIR4RS principles^41,42^ (Table S4). Community participation is welcomed, ensuring that FERMO remains a robust, evolving tool for metabolomics research.

FERMO also has certain limitations, primarily related to the type of input data it supports. It is specifically designed to process data from liquid chromatography electrospray ionization (tandem) mass spectrometry (LC-ESI-MS/MS) run in untargeted data-dependent acquisition (DDA) mode of a single polarity (either positive or negative ion mode), with samples acquired at identical concentration, dilution, and injection volume. Additionally, FERMO requires a pre-formatted peak table rather than raw LC-MS/MS data. While this design choice simplifies data handling and reduces infrastructure demands (peak table files are significantly smaller in size than raw data files), it also necessitates that users preprocess their data before analysis, adding an extra layer of complexity to the workflow. Importantly, FERMO requires the feature-width-at-half-maximum (FWHM) to construct its pseudo-chromatogram visualization. To our knowledge, the tool mzmine (versions 3 and 4) is currently the only GUI tool that reports this value natively, and therefore, FERMO presently only accepts such formatted peak tables. We encourage developers of other metabolomics processing tools to include the FWHM in their output to allow compatibility with FERMO.

FERMO’s prioritization framework is mostly qualitative by nature, focusing on hypothesis-driven selection rather than statistical analysis of metabolomics datasets. Yet, compared to manual exploration and interpretation of spectral data, it facilitates prioritization based on clearly defined quantitative and reproducible scores, making the process more objective and standardized. We note that FERMO prioritizes “by elimination” mostly, leaving the metabolite features that are most likely associated with the research question represented by the set filters. Researchers requiring advanced statistical interpretation may find dedicated tools such as MetaboAnalyst^15^ more suitable for their needs. Here, we also stress that correlation does not equal causation and that any prioritized samples and metabolite features should be validated in follow-up experiments.

Since FERMO’s visualization is browser-based, large-scale analyses may exceed browser-specific display limitations, although exploratory benchmarking showed that jobs with ∼3600 features were still tolerated (Table S2). For such cases, users are provided with a download page where session files and spectral networks can be accessed for visualization in external tools like Cytoscape. Moving forward, we plan to optimize FERMO’s performance, improve processing speed, and expand data visualization capabilities. We aim to integrate additional input formats, accommodate more types of orthogonal data, and refine the user experience. To ensure that FERMO continues to evolve in alignment with community needs, we encourage user feedback and collaboration through the FERMO Discussions board (https://github.com/orgs/fermo-metabolomics/discussions).

In conclusion, the FERMO dashboard provides a robust and versatile framework for sample and feature prioritization based on generalized principles. By combining intuitive visualization with a range of attribute filters, FERMO enables rapid and streamlined data exploration, as demonstrated in the case studies. Its ability to integrate and organize LC-MS/MS data in a centralized, accessible manner makes it a valuable tool for hypothesis-driven metabolomics research. We anticipate that FERMO will support diverse applications across the life sciences, facilitating data-driven decision-making in fields such as natural product discovery, environmental metabolomics, and biomedical research.

## Methods

### Software design overview

FERMO is an open-source GUI application designed for hypothesis-driven prioritization of metabolomics data, available at https://fermo.bioinformatics.nl/. FERMO is free to users from both academia and industry, requires no registration, and does not limit the number of analysis runs. Results are stored for a grace period of 30 days, during which they can be freely shared and downloaded via their unique FERMO job ID. Of note, FERMO does not guarantee the confidentiality of uploaded data, although access of results without possession of the FERMO job ID is highly unlikely. Briefly, the software is structured into two interconnected modules. FERMO GUI (presently, version 1.1.2) is the browser-based frontend for user interaction, job submission, and result visualization. Although primarily designed as a web application, the FERMO GUI is also available as a Docker container, enabling local installation. Fermo_core (presently, version 0.6.4) is the backend module responsible for executing analyses and data processing. Fermo_core can also be used as a stand-alone CLI tool, offering flexibility for advanced users, allowing integration into custom workflows, and the analyses of larger datasets without the constraints imposed by direct visualization within an interactive dashboard. Both modules are implemented in Python 3.11, adhere to semantic versioning, and follow a modular design philosophy to ensure scalability and maintainability. For a detailed description of the software architecture and its modules, see the Supplementary Methods or the online documentation at https://fermo-metabolomics.github.io/fermo_docs/.

### General information on statistical methods

To identify phenotype-associated features, FERMO applies two-sided Pearson correlation using the scipy.stats.pearsonr() function (Python library SciPy v1.10.1). Correlations are calculated between z-transformed feature intensities and z-transformed quantitative phenotype values, limited to features detected in at least four samples. The resulting p-values are subjected to FDR correction using a user-specified method selected from statsmodels.stats.multitest.multipletests() (Statsmodels v0.14.4). In the *“Benchmarking FERMO against established phenotype prioritization workflows”* (14 samples, no replicates), Benjamini-Hochberg correction was applied. In *“FERMO effectively prioritizes bioactive actinomycin D in an OSMAC study”* (36 samples, no replicates), Bonferroni correction was used. Both corrected and uncorrected p-values are reported alongside the corresponding Pearson correlation coefficients (*r*) as effect size estimates. Of note, users of FERMO are solely responsible for the experimental design, including the number of samples and replicates, and thus the statistical power of their experiments.

### Benchmarking FERMO against established phenotype prioritization workflows: Bioactivity-Based Molecular Networking & NP3 MS Workflow

#### Data retrieval and preprocessing

LC-MS/MS data in .mzML format of full extract and fractions used by the Bioactivity-Based Molecular Networking study was downloaded from the MassIVE repository (MassIVE ID MSV000080502). Sample blank measurements were excluded from the analysis, resulting in a total of 14 files processed in the analysis. Selectivity indices for extracts and fractions were collected from the original publication and converted into the FERMO-compatible “percentage” format (higher selectivity indices signify more potent values). LC-MS/MS data was pre-processed using mzmine3 version 3.9.0 in batch mode, using the batch mode .xml-file (Table S1F). Briefly, mass ion peaks were detected for MS1 and MS2 at a noise level of 1.0E3 and 1.0E2, respectively, (positive polarity, mass detector: centroid) and their chromatograms were built using ADAP chromatogram builder (retention time cutoff 5-38 mins, minimum group size in number of scans: 8; min intensity scans: 1.0E3; group intensity threshold: 5.0E3). The chromatograms were deconvoluted (algorithm: local minimum feature resolver; chromatographic threshold: 90; search minimum in RT range: 0.2; minimum relative height: 0%; minimum ratio of peak top/edge: 2.0; peak duration: 0.01-2 min, minimum scans: 6). The detected peaks were deisotoped (maximum charge: 3; retention time tolerance: 0.1 min; monotonic shape; representative isotope: lowest *m/z*). Peak lists from different extracts were aligned (algorithm: join aligner; weight for RT = 1; RT tolerance: 0.3; weight for *m/z* = 3). Missing peaks detected in at least one of the samples were filled with the peak finder algorithm (Intensity tolerance: 10%; RT tolerance: 0.4 min, minimum scans: 1). Peaks were filtered using the Feature list row filter and only retained if detected in at least 2 samples and having a feature duration range of 0.3-3.0 mins and a chromatographic FWHM of 0.15-1.0. All modules used 15 ppm as mass deviation tolerance. The resulting peak table contained 912 features and was exported for consecutive processing using the Molecular Networking File Export module (Filter rows: ALL; Feature intensity: Height; CSV export: COMPREHENSIVE). The resulting .mgf-file containing MS/MS spectra was analysed using MS2Query version 1.5.3 in positive mode, using the MS2Query library files v8 available from Zenodo^43^.

#### FERMO parameters and analysis settings

Analysis using the mzmine3-generated peak table, MS/MS information, phenotype data, and MS2Query annotations was performed with fermo_core 0.6.2 in command line mode. Detailed file and parameter settings are referenced in Table S1G. Briefly, feature and sample information was parsed from the file case1_quant_full.csv in the format ‘mzmine3’ with positive ion mode polarity. MS/MS fragmentation information was parsed from the file case1.mgf in the format ‘mgf’. MS/MS fragments +- 10 mass units around precursor *m/z* were removed. Phenotype/bioactivity information was parsed from the file case1_phenotype.csv in the format ‘quantitative-percentage’. MS2Query results were parsed from the file case1_ms2query.csv. Only results with a score above a user-specified value of 0.7 were retained.

Molecular features were filtered and only retained if they were inside the relative area range of 0.01-1.0 in at least one sample (relative to the feature with the highest area in the sample). For each sample, overlapping features were annotated for ion adducts using a cutoff mass deviation value of 15.0 ppm. MS/MS spectra of all features with more than 12 fragment ions were compared pairwise and scored using the ‘modified cosine’ algorithm, with a fragment tolerance of 0.1. From the resulting similarity matrix, a network was created, with features represented as nodes and the similarity value as edges. Edges were pruned if their score was below a similarity cutoff of 0.7. Also, edges were pruned so that only the 10 highest scoring edges remained. MS/MS spectra of all features with more than 12 fragment ions were compared pairwise and scored using the MS2Deepscore algorithm. From the resulting similarity matrix, a network was created, with features represented as nodes and the similarity value as edges. Edges were pruned if their score was below a similarity cutoff of 0.8. Also, edges were pruned so that only the 10 highest scoring edges remained. For each feature detected in more than three phenotype-associated samples, the mean area was correlated with the percentage activity per sample using Pearson correlation and the feature was only considered phenotype-associated if its coefficient was greater than 0.8 and its FDR-corrected p-value (Benjamini Hochberg method) less than 0.05. The resulting out.fermo.session.json file was visualized with FERMO GUI v1.1.0 and is referenced in the Data Availability Statement and Supplementary Table S1G.

#### Data analysis with NP3 MS workflow

NP3 MS Workflow (v1.1.6) was downloaded and installed following instructions on its GitHub page. LC-MS/MS data in .mzML format of full extract and fractions used by the Bioactivity-Based Molecular Networking was downloaded from MSV000080502. Sample blank measurements were excluded from the analysis, resulting in a total of 14 files processed in the analysis. Selectivity indices for extracts and fractions were collected from the original BioMN publication and scaled between 0 and 100, following instructions in the NP3 MS Workflow documentation. A metadata file was constructed using the provided metadata template and NP3 MS Workflow was executed with default settings, using the “run” command.

### FERMO effectively prioritizes bioactive actinomycin D in an OSMAC study

#### Bacterial strains, growth conditions and metabolite extraction

*Streptomyces* sp. MBT27, previously isolated from the Qingling Mountains, Shanxi province, China, was obtained from the Leiden University strain collection^44^. *Streptomyces* sp. MBT27 was grown in liquid Minimal Medium supplemented with various carbon sources as previously described^34^. The carbon sources (percentages in w/v) were: 1% mannitol + 1% glycerol, 1% mannitol, 2% mannitol, 1% glycerol, 2% glycerol, 1% glucose, 2% glucose, 1% fructose, 1% arabinose or 1% N-acetylglucosamine (GlcNAc). After incubation, culture supernatants were extracted with ethyl acetate (EtOAc).

#### Antimicrobial activity assay

The antimicrobial activity of the crude extract was evaluated using a disc diffusion assay, targeting methicillin-sensitive *Staphylococcus aureus* ATCC29213. For this assay, 40 µg of the crude extract was dissolved in MeOH and applied onto sterile discs, which were then dried and placed on agar plates pre-inoculated with the bacterial strain. After incubation, the size of the inhibition halos around the discs were measured and converted to percentage relative to the largest growth inhibition halo. Ampicillin and MeOH served as the positive and negative controls, respectively.

#### Data-dependent LC-ESI-HRMS/MS

LC-MS/MS acquisition was performed using a Shimadzu Nexera X2 ultra-high-performance liquid chromatography system, with an attached photodiode array detector, coupled to Shimadzu 9030 QTOF mass spectrometer, equipped with a standard electrospray ionization source unit with a calibrant delivery system, as described previously^45,46^. A total of 1 µL was injected into a Waters Acquity HSS C18 column (1.8 μm, 100 Å, 2.1 × 100 mm). The column was maintained at 30 °C, and run at a flow rate of 0.5 mL/min, using 0.1% formic acid in H2O (solvent A) and 0.1% formic acid in acetonitrile (solvent B). The gradient used was 5% B for 1 min, 5-85% B for 9 min, 85-100% B for 1 min, and 100% B for 4 min. The column was re-equilibrated to 5% B for 3 min before the next run was started. The PDA acquisition was performed in the range of 200-600 nm, at 4.2 Hz, with a 1.2 nm slit width. The flow cell was maintained at 40 °C. All the samples were analyzed in positive ion polarity, using data-dependent acquisition mode. In this regard, full scan MS spectra (*m/z* 100-1700, scan rate 10 Hz, ID enabled) were followed by two data-dependent MS/MS spectra (*m/z* 100-1700, scan rate 10 Hz, ID disabled) for the two most intense ions per scan. The ions were fragmented using collision-induced dissociation (CID) with fixed collision energy (CE 20 eV) and excluded for 1 s before being re-selected for fragmentation. The parameters used for the ESI source were: interface voltage 4 kV, interface temperature 300 °C, nebulizing gas flow 3 L/min, and drying gas flow 10 L/min.

#### Data processing: mzmine3 pre-processing parameters

LC-MS/MS data was converted using Shimadzu LabSolutions Postrun Analysis to the .mzXML format, and was processed by mzmine 3.3.0. Briefly, mass ion peaks were detected for MS1 and MS2 at a noise level of 1.0E3 and 1.0E1, respectively, (positive polarity, mass detector: centroid) and their chromatograms were built using ADAP chromatogram builder (minimum group size in number of scans: 10; group intensity threshold: 3.0E3). The chromatograms were deconvoluted (algorithm: local minimum feature resolver; chromatographic threshold: 90; search minimum in RT range: 0.05; minimum relative height: 0%; minimum ratio of peak top/edge: 1.8; peak duration: 0.01-1.5 min). The detected peaks were deisotoped (maximum charge: 3; representative isotope: most intense). Peak lists from different samples were aligned (weight for RT =1; weight for *m/z* = 4; mobility weight 1.0). Features detected in at least one of the samples were gap-filled with the peak finding algorithm (RT tolerance: 0.1 min, *m/z* tolerance was set to 0.002 *m/z* or 10 ppm). Duplicate peaks were filtered. Only the features with MS/MS data were exported for consecutive processing using the Molecular networking File export module (Filter rows: ALL; Feature intensity: Height; CSV export: COMPREHENSIVE). The resulting .mgf-file containing MS/MS spectra was analysed using MS2Query version 1.5.3 in positive mode, using the MS2Query library files v8 available from Zenodo^43^.

#### FERMO parameters and analysis settings

Analysis using the mzmine3-generated peak table, MS/MS information, phenotype data, sample group metadata, MS2Query annotations, and antiSMASH KnownClusterBlast output was performed with fermo_core 0.6.2 in command line mode. Detailed file and parameter settings are referenced in Table S1H. Briefly, feature and sample information was parsed from the file 20250310_all_CS_quant_full.csv in the format ‘mzmine3’ with positive ion mode polarity. MS/MS fragmentation information was parsed from the file 20250310_all_CS.mgf in the format ‘mgf’. MS/MS fragments +- 10 mass units around precursor *m/z* were removed. MS/MS fragments with an intensity less than 0.01 relative to the base peak were removed. Phenotype/bioactivity information was parsed from the file bioactivity_all_cs_perc_correct.csv in the format ‘quantitative-percentage’. Group metadata information was parsed from the file metadataFERMO_all_cs.csv in the format ‘fermo’. MS2Query results were parsed from the file 20250310_all_CS_ms2query.csv. Only results with a score above a user-specified value of 0.7 were retained. antiSMASH results were parsed from the directory NZ_CP070373.1. KnownClusterBlast matches were only retained if they were above a user-specified similarity cutoff of 0.7. For each sample, overlapping features were annotated for ion adducts using a cutoff mass deviation value of 10.0 ppm. For each feature, neutral losses were calculated between the precursor *m/z* and each MS/MS fragment peak *m/z* and matched against a library of annotated neutral losses, using a cutoff mass deviation value of 10.0 ppm. For each feature, MS/MS fragments were matched against a library of annotated MS/MS fragments, using a cutoff mass deviation value of 10.0 ppm. MS/MS spectra of all features with more than 5 fragment ions were compared pairwise and scored using the ‘modified cosine’ algorithm, with a fragment tolerance of 0.1. From the resulting similarity matrix, a network was created, with features represented as nodes and the similarity value as edges. Edges were pruned if their score was below a similarity cutoff of 0.8. Also, edges were pruned so that only the 5 highest scoring edges remained. MS/MS spectra of all features with more than 5 fragment ions were compared pairwise and scored using the MS2Deepscore algorithm. From the resulting similarity matrix, a network was created, with features represented as nodes and the similarity value as edges. Edges were pruned if their score was below a similarity cutoff of 0.8. Also, edges were pruned so that only the 10 highest scoring edges remained. Features only detected in sample-blanks were considered blank-associated, as were features that had a quotient of less than 10 when their mean area between samples and sample blanks was compared. Samples were grouped according to the provided group metadata information. For each feature observed in more than one group, the quotient between the mean area of groups was calculated pairwise. For each feature detected in more than three phenotype-associated samples, the mean area was correlated with the percentage activity per sample using Pearson correlation and the feature was only considered phenotype-associated if its coefficient was greater than 0.7 and its Bonferroni-corrected *p*-value less than 0.05. The MS/MS spectrum of each feature was matched pairwise against a targeted spectral library constructed from relevant matches of the KnownClusterBlast algorithm. Matching was performed using the ‘modified cosine’ algorithm, with a fragment tolerance of 0.1 and matches were only retained if the number of matched peaks between feature and library spectrum was greater than 5, the score exceeded the cutoff score of 0.7, and the maximum precursor *m/z* difference of 600 was not exceeded. The resulting out.fermo.session.json file was visualized with FERMO GUI v1.1.0 and is referenced in Table S1H.

## Data Availability Statement

The dataset generated and analyzed in section *FERMO effectively prioritizes bioactive actinomycin in an OSMAC study* was deposited on MassIVE under MSV000097564 available at https://doi.org/doi:10.25345/C5JS9HM3J^47^. All results generated by FERMO and discussed through the manuscript are available at https://doi.org/10.5281/zenodo.15203137^48^ (*Benchmarking FERMO against established phenotype-prioritization workflows* contained in directory “case_study_1”, *FERMO effectively prioritizes bioactive actinomycin D in an OSMAC study* in “case_study_2”).

## Code Availability Statements

The FERMO dashboard is freely available at https://fermo.bioinformatics.nl/ with no login requirements or job quota limits, and is expected to remain accessible for the foreseeable future. The source code for the FERMO dashboard is openly available at GitHub (https://github.com/fermo-metabolomics/fermo) and archived on Zenodo (https://doi.org/10.5281/zenodo.7565700^49^). The source code for the fermo_core CLI is available at GitHub (https://github.com/fermo-metabolomics/fermo_core) and archived on Zenodo (https://doi.org/10.5281/zenodo.11259126^50^). All code is licensed under the Open Source Initiative-approved MIT License, permitting unrestricted academic and commercial use.

## Acknowledgements

The authors thank Kevin Mildau, Luis Manuel Quirós-Guerrero, Louis-Félix Nothias, Adriano Rutz, Jean-Luc Wolfender, and participants of the 2022 Dagstuhl Conference with the title: “Computational Metabolomics: From Spectra to Knowledge” for valuable inspiration. The authors acknowledge the project contribution of Koen van Ingen. The authors are thankful to the beta version testers for their effort and feedback (in alphabetical order): Esteban Charria Girón, Maria Dell, Marianna Iorio, Soliman Khatib, George Lund, Sonia Maffioli, Leo Padva, and Matteo Simone. This project has received funding from the European Union’s Horizon 2020 research and innovation programme under Grant Agreement no. 101000392 (MARBLES) and by the NWO Grant KICH1.LWV04.21.013.

## Author Contributions

M.M.Z., M.H.M, and J.J.J.v.d.H conceptualized the project idea. M.M.Z. contributed to all parts of the project and designed and programmed the software backend and frontend, performed data analysis and benchmarking, and wrote the original manuscript. H.E.A. contributed to dashboard design, programming, and original manuscript; N.V.M. and L.M.B. performed experimental investigation and data analysis. S.S., N.d.J., and S.C. contributed to software programming and/or testing. G.P.v.W. revised the manuscript and provided project supervision. M.H.M. and J.J.J.v.d.H. contributed to software design, revised the manuscript, and provided project supervision. All co-authors have read and agree with the content of the manuscript.

## Materials & Correspondence

Please direct all questions regarding computer code to M.M.Z. and/or post them in the FERMO Metabolomics Discussion Forum (https://github.com/orgs/fermo-metabolomics/discussions).

## References

1. Eggert, U. S. The why and how of phenotypic small-molecule screens. Nat Chem Biol 9, 206–209 (2013).

2. Niessen, W. M. A. & Correa C. R. A. Interpretation of MS-MS Mass Spectra of Drugs and Pesticides. (John Wiley & Sons, 2017).

3. Stone, M. J. & Williams, D. H. On the evolution of functional secondary metabolites (natural products). Mol Microbiol 6, 29–34 (1992).

4. Newman, D. J. & Cragg, G. M. Natural Products as Sources of New Drugs over the Nearly Four Decades from 01/1981 to 09/2019. J Nat Prod 83, 770–803 (2020).

5. Wolfender, J.-L., Nuzillard, J.-M., van der Hooft, J. J. J., Renault, J.-H. & Bertrand, S. Accelerating Metabolite Identification in Natural Product Research: Toward an Ideal Combination of Liquid Chromatography-High-Resolution Tandem Mass Spectrometry and NMR Profiling, in Silico Databases, and Chemometrics. Anal Chem 91, 704–742 (2019).

6. Quinn, R. A. et al.. Molecular Networking As a Drug Discovery, Drug Metabolism, and Precision Medicine Strategy. Trends Pharmacol Sci 38, 143–154 (2017).

7. Beniddir, M. A. et al.. Advances in decomposing complex metabolite mixtures using substructure- and network-based computational metabolomics approaches. Nat Prod Rep 38, 1967–1993 (2021).

8. Smith, C. A., Want, E. J., O’Maille, G., Abagyan, R. & Siuzdak, G. XCMS: processing mass spectrometry data for metabolite profiling using nonlinear peak alignment, matching, and identification. Anal Chem 78, 779–787 (2006).

9. Schmid, R. et al.. Integrative analysis of multimodal mass spectrometry data in MZmine 3. Nat Biotechnol 41, 447–449 (2023).

10. Pfeuffer, J. et al.. OpenMS - A platform for reproducible analysis of mass spectrometry data. J Biotechnol 261, 142–148 (2017).

11. Wang, M. et al.. Sharing and community curation of mass spectrometry data with Global Natural Products Social Molecular Networking. Nat Biotechnol 34, 828–837 (2016).

12. Takeda, H. et al.. MS-DIAL 5 multimodal mass spectrometry data mining unveils lipidome complexities. Nat Commun 15, 9903 (2024).

13. Dührkop, K. et al.. SIRIUS 4: a rapid tool for turning tandem mass spectra into metabolite structure information. Nat Methods 16, 299–302 (2019).

14. Letourneau, D. R. & Volmer, D. A. Constellation: An Open-Source Web Application for Unsupervised Systematic Trend Detection in High-Resolution Mass Spectrometry Data. J Am Soc Mass Spectrom 33, 382–389 (2022).

15. Pang, Z. et al.. MetaboAnalyst 6.0: towards a unified platform for metabolomics data processing, analysis and interpretation. Nucleic Acids Res 52, W398–W406 (2024).

16. Mildau, K. et al.. Effective data visualization strategies in untargeted metabolomics. Nat Prod Rep (2024) doi:10.1039/d4np00039k.

17. Samples, R. M., Puckett, S. P. & Balunas, M. J. Metabolomics Peak Analysis Computational Tool (MPACT): An Advanced Informatics Tool for Metabolomics and Data Visualization of Molecules from Complex Biological Samples. Anal Chem 95, 8770–8779 (2023).

18. Covington, B. C. & Seyedsayamdost, M. R. MetEx, a Metabolomics Explorer Application for Natural Product Discovery. ACS Chem Biol 16, 2825–2833 (2021).

19. Nothias, L.-F. et al.. Bioactivity-Based Molecular Networking for the Discovery of Drug Leads in Natural Product Bioassay-Guided Fractionation. J Nat Prod 81, 758–767 (2018).

20. Olivon, F. et al.. Bioactive Natural Products Prioritization Using Massive Multi-informational Molecular Networks. ACS Chem Biol 12, 2644–2651 (2017).

21. Ory, L. et al.. Targeting bioactive compounds in natural extracts - Development of a comprehensive workflow combining chemical and biological data. Anal Chim Acta 1070, 29–42 (2019).

22. Bazzano, C. F. et al.. NP MS Workflow: An Open-Source Software System to Empower Natural Product-Based Drug Discovery Using Untargeted Metabolomics. Anal Chem 96, 7460–7469 (2024).

23. Lee, S. et al.. NP Analyst: An Open Online Platform for Compound Activity Mapping. ACS Cent Sci 8, 223–234 (2022).

24. Zdouc, M. M. et al.. FERMO: A dashboard for streamlined rationalized prioritization of molecular features from mass spectrometry data. bioRxiv (2022) doi:10.1101/2022.12.21.521422.

25. Tabudravu, J. N. et al.. LC-HRMS-Database Screening Metrics for Rapid Prioritization of Samples to Accelerate the Discovery of Structurally New Natural Products. J Nat Prod 82, 211–220 (2019).

26. Pérez-Victoria, I., Martín, J. & Reyes, F. Combined LC/UV/MS and NMR Strategies for the Dereplication of Marine Natural Products. Planta Med 82, 857–871 (2016).

27. Zdouc, M. M. et al.. Planomonospora: A Metabolomics Perspective on an Underexplored Actinobacteria Genus. J Nat Prod 84, 204–219 (2021).

28. Zdouc, M. M. et al.. A biaryl-linked tripeptide from Planomonospora reveals a widespread class of minimal RiPP gene clusters. Cell Chem Biol 28, 733–739.e4 (2021).

29. Shannon, P. et al.. Cytoscape: a software environment for integrated models of biomolecular interaction networks. Genome Res 13, 2498–2504 (2003).

30. de Jonge, N. F. et al.. MS2Query: reliable and scalable MS mass spectra-based analogue search. Nat Commun 14, 1752 (2023).

31. Blin, K. et al.. antiSMASH 8.0: extended gene cluster detection capabilities and analyses of chemistry, enzymology, and regulation. Nucleic Acids Res (2025) doi:10.1093/nar/gkaf334.

32. Heuckeroth, S. et al.. Reproducible mass spectrometry data processing and compound annotation in MZmine 3. Nat Protoc 19, 2597–2641 (2024).

33. Bode, H. B., Bethe, B., Höfs, R. & Zeeck, A. Big effects from small changes: possible ways to explore nature’s chemical diversity. Chembiochem 3, 619–627 (2002).

34. Machushynets, N. V. et al.. Discovery of actinomycin L, a new member of the actinomycin family of antibiotics. Sci Rep 12, 2813 (2022).

35. Zdouc, M. M. et al.. MIBiG 4.0: advancing biosynthetic gene cluster curation through global collaboration. Nucleic Acids Res 53, D678–D690 (2025).

36. Rutz, A. et al.. Taxonomically Informed Scoring Enhances Confidence in Natural Products Annotation. Front Plant Sci 10, 1329 (2019).

37. Petras, D. et al.. GNPS Dashboard: collaborative exploration of mass spectrometry data in the web browser. Nat Methods 19, 134–136 (2022).

38. Mejri, Y. et al.. MS2DECIDE: Aggregating multi-annotated tandem mass spectrometry data with Decision Theory enhances natural products prioritization. ChemRxiv (2024) doi:10.26434/chemrxiv-2024-x6p8r.

39. Mildau, K. et al.. Tailored Mass Spectral Data Exploration Using the SpecXplore Interactive Dashboard. Anal Chem 96, 5798–5806 (2024).

40. Quiros-Guerrero, L.-M. et al.. INVENTA: A computational tool to discover structural novelty in natural extracts libraries. Front Mol Biosci 9, 1028334 (2022).

41. Hoyt, C. T. & Gyori, B. M. The O3 guidelines: open data, open code, and open infrastructure for sustainable curated scientific resources. Sci Data 11, 547 (2024).

42. Barker, M. et al.. Introducing the FAIR Principles for research software. Sci Data 9, 622 (2022).

43. de Jonge, N. MS2Query pre-trained embeddings and models. Zenodo 10.5281/ZENODO.13348638 (2024).

44. Zhu, H. et al.. Eliciting antibiotics active against the ESKAPE pathogens in a collection of actinomycetes isolated from mountain soils. Microbiology (Reading) 160, 1714–1725 (2014).

45. Machushynets, N. V. et al.. Discovery and Derivatization of Tridecaptin Antibiotics with Altered Host Specificity and Enhanced Bioactivity. ACS Chem Biol 19, 1106–1115 (2024).

46. Machushynets, N. V. et al.. Exploring the Chemical Space of NRPs and Discovery of Paenilipoheptin B. Org Lett 27, 2821–2825 (2025).

47. van Wezel, G. P. MassIVE MSV000097564 - OSMAC Study on Streptomyces sp MBT27. MassIVE 10.25345/C5JS9HM3J (2025).

48. Zdouc, M. M. Fermo-Metabolomics/fermo_ms Zenodo doi:10.5281/ZENODO.15203137 (2025).

49. Zdouc, M. M., de Jonge, N. & Augustijn, FERMO. Zenodo doi:10.5281/ZENODO.7565700 (2024).

50. Zdouc, M. M. Fermo_core. Zenodo. doi:10.5281/ZENODO.11259126 (2025).

