## Supplemental Information for "FERMO: a Dashboard for Automated Prioritization of Molecular Features from Mass Spectral Data"

### Table of contents

**FERMO: a Dashboard for Automated Prioritization of Molecular Features from Mass Spectral Data - Supplementary Information** 1
Supplementary Methods 3
FERMO software design overview 3
System Architecture Overview 3
Frontend organization 3
Backend organization 3
Data Handling & Formats 4
Availability and Installation 5
Data processing 5
Filter modules 5
Network modules 5
Annotation modules 5
Sample group assignment 6
Phenotype data assignment 6
Qualitative Data (Binary: Positive/Negative) 6
Quantitative Data – Percentage-Based Measurements 7
Quantitative Data – Concentration-Based Measurements 7
Scores calculation 8
Feature Phenotype Score 8
Feature Novelty Score 8
Sample Mean Novelty Score 8
Sample Diversity Score 8
Sample Specificity Score 8
Supplementary Data 10
Supplementary Table S1: Collection of hyperlinks referenced in main text 10
Supplementary Table S2: Benchmarking of FERMO run time 10
Supplementary Table S3: Qualitative comparison of metabolomics prioritization tools with FERMO 12
Supplementary Table S4: FAIR4RS Checklist 13
Supplementary Figure S1: Comparison pseudo-chromatograms with total ion chromatograms 14
References 15

### Supplementary Methods

#### FERMO software design overview

##### System Architecture Overview

FERMO is designed with a client-server architecture, where the frontend and backend communicate via standardized JSON-based requests. The system is built for modularity (components can be independently updated or extended), scalability (small and large metabolomics datasets can be handled efficiently), and reproducibility (registering parameters, processing steps, and version-controlled dependencies).

##### Frontend organization

FERMO is implemented as a Flask v3 application. The frontend is responsible for i) **user input handling,** accepting user-provided files and parameters while ensuring data validation; ii) **job management,** dispatching analysis requests to the backend using a structured parameter file; and iii) **result presentation,** displaying analysis outputs via an interactive dashboard. To handle asynchronous task execution, FERMO integrates Celery v5 as a task queue manager, with Redis serving as a message broker. The system is optimized for easy deployment using Docker v3.3, allowing platform-independent installation. Additionally, FERMO can be installed locally as described in section Availability and Installation. A detailed list of dependencies is available at https://github.com/fermo-metabolomics/FERMO/blob/main/fermo_gui/pyproject.toml.

##### Backend organization

The backend, fermo_core, is responsible for executing analysis workflows. It operates as a stand-alone command line interface (CLI) or as a backend service for the GUI. The backend follows a structured pipeline consisting of i) **input handling,** parsing and validating user-submitted parameter files (JSON format, controlled via JSON Schema); ii) **data processing,** applying analysis steps described in detail below; and iii) **output generation,** producing standardized output files to be processed by FERMO GUI or other downstream tools. Since the CLI version of fermo_core requires manual creation of a parameter file following the provided JSON Schema specifications, its use is only recommended for advanced users comfortable with command-line operations. To ensure robustness and reliability, fermo_core follows test-driven development[1], with 90% test coverage across 558 unit and integration tests (as of version 0.6.1), automated testing via continuous integration, and structured error handling to minimize failures during long-running analysis jobs. A detailed list of dependencies is available at https://github.com/fermo-metabolomics/fermo_core/blob/main/pyproject.toml.

#### Data Handling & Formats

FERMO processes metabolomics data acquired through liquid chromatography-electrospray ionization (LC-ESI) data-dependent acquisition (DDA) tandem mass spectrometry (MS/MS). To ensure data consistency and reliability, input files must contain measurements from a single polarity mode, either positive or negative, without mixing. All samples should be acquired under identical experimental conditions, including the same concentration or dilution and injection volume, to minimize variability. Additionally, high-resolution mass spectrometry data with a mass accuracy of ≤ 20 ppm is recommended to improve feature identification and reduce mass deviation errors. Input data formats accepted by and output data produced by FERMO are summarized in the tables below. Detailed information is available from the Documentation at https://fermo-metabolomics.github.io/fermo_docs/home/input_output/.

**Overview of data types and formats accepted by FERMO**

| Data type | Format | Mandatory/Optional |
|---|---|---|
| Molecular feature peak table | Mzmine3, Mzmine4 | mandatory |
| MS/MS spectrum information | mgf (Mzmine3, Mzmine4) | optional |
| Sample grouping metadata | FERMO | optional |
| Phenotype data | FERMO (qualitative, quantitative–percentage, quantitative-concentration) | optional |
| Spectral library | mgf | optional |
| MS2Query results file | MS2Query* | optional |
| antiSMASH results | antiSMASH job ID | optional |

*needs an additional column “id” or “feature_id” corresponding to peak table (by default contained in output of MS2Query >= v1.5.3)

**Overview of output generated by FERMO**

| Suffix | Description |
|---|---|
| .session.json | Complete FERMO analysis results, used as input for dashboard. |
| .summary.txt | Human-friendly summary of processing steps performed. |
| .graphml | Cytoscape-compatible spectral similarity (=molecular) network file, one per algorithm applied |
| .fermo.abbrev.csv | Cytoscape-compatible feature annotation file. |
| .fermo.full.csv | Input peak table modified by annotations generated by FERMO, usable in downstream processing. |
| .log | Full log of all processing steps performed, including warnings and/or errors. |

##### Availability and Installation

FERMO is available as a web application at https://fermo.bioinformatics.nl/ with certain usage restrictions to prevent excessive strain on computational resources, including a file size limit of 9 MB, a maximum runtime of 60 min, and job storage of 30 days. For unrestricted use, FERMO can be installed locally, with source code available at https://github.com/fermo-metabolomics/FERMO. The standalone fermo_core CLI is distributed as a Python package and can be installed directly from PyPI at https://pypi.org/project/fermo-core/ or from source code available at https://github.com/fermo-metabolomics/fermo_core.

#### Data processing

FERMO organizes data processing into a modular framework, allowing flexibility in analysis workflows. By default, a recommended set of modules and parameters is preconfigured to facilitate a streamlined "quickstart" process. However, all modules and parameters can be modified, disabled, or customized, and additional modules can be enabled as needed to accommodate specific research requirements.

##### Filter modules

FERMO includes filtering steps for both molecular features and MS/MS spectra to reduce the overall complexity of the analysis. Molecular feature filtering allows the retention of only those features that exceed a specified relative intensity or area in at least one sample. MS/MS spectra can be refined by removing the precursor *m/z* along with all fragment ions within ±10 *m/z* units. Additionally, fragment ions with a relative intensity below a user-defined threshold can be excluded to improve downstream analysis.

##### Network modules

FERMO organizes molecular features using spectral similarity networking, also referred to as molecular networking[2]. In this approach, MS/MS fragmentation spectra are compared in a pairwise manner, and features exhibiting similar spectra are grouped together. This method is based on the principle that structurally related molecules typically produce similar fragmentation patterns, allowing for the identification of chemical families. FERMO supports two spectral similarity algorithms in parallel: the modified cosine similarity[3] and MS2DeepScore[4]. The resulting network-based relationships among molecular features are subsequently used to compute Sample Diversity and Specificity Scores, as described in the following sections.

##### Annotation modules

FERMO annotates molecular features using a multi-step approach that integrates ion identity assignment, fragment annotation, spectral library matching, and genome-based molecular feature association. First, FERMO identifies co-occurring molecular features, those with overlapping retention time ranges, and assigns their ion identities for both positive and negative ion modes, covering 20 and 6 types of adducts, respectively (for a complete list, see the FERMO Documentation at https://fermo-metabolomics.github.io/fermo_docs/modules/annotation.adduct/). Next, MS/MS fragment ions (such as $y_2$- and $b_2$-fragments from proteinogenic dipeptides) and neutral losses (calculated from precursor *m/z*) are annotated using literature-reported databases[5–7]. Spectral library matching is performed against one or more user-provided libraries using the Modified Cosine[3] and MS2DeepScore[4] spectral similarity algorithms. Notably, this process can be computationally intensive, so the use of targeted libraries is recommended to optimize performance. Alternatively, FERMO supports results generated by MS2Query[8], a highly efficient annotation tool whose output can be directly integrated into FERMO. Additionally, FERMO accepts functional genomics annotations generated by the antiSMASH web server (versions 7 and 8)[9]. In particular, FERMO parses results from antiSMASH's KnownClusterBlast module, which compares the sequence similarity of biosynthetic gene clusters annotated by antiSMASH against literature-reported BGCs deposited in the Minimum Information about a Biosynthetic Gene Cluster (MIBiG) database [10]. FERMO parses the KnownClusterBlast results, selects matches exceeding a user-defined similarity threshold, and uses it to subset the GNPS spectral library[11] and an *in silico* spectral library calculated from structures in MIBiG 3.0[12] using the tool CFM-ID 4.0[13]. This targeted spectral library is annotated with MIBiG IDs, allowing linking of matched molecular features to BGCs putatively responsible for their production. Again, spectral matching can be performed using the Modified Cosine[3] and MS2DeepScore[4] spectral similarity algorithms.

#### Sample group assignment

FERMO annotates molecular features with sample grouping information to assess intergroup variability. Sample blanks are designated separately to distinguish background signals from biologically relevant features. To determine blank association, the mean, median, or maximum area or height of each molecular feature is calculated for both blank and non-blank samples. The quotient of these values is then compared to a user-defined fold-change threshold. If the quotient exceeds this threshold, the feature is considered not blank-associated and is retained for further analysis. This approach helps to distinguish features arising from sample cross-contamination due to column retention/bleed.

#### Phenotype data assignment

FERMO accepts both qualitative and quantitative phenotype data, processing each input format with a dedicated algorithm. Input files must follow a certain format as described in detail in the FERMO documentation at https://fermo-metabolomics.github.io/fermo_docs/home/input_output/.

##### Qualitative Data (Binary: Positive/Negative)

FERMO accepts qualitative phenotype data in binary format (e.g. positive/negative, active/inactive, presence/absence, ect). The algorithm for qualitative phenotype data functions as follows: Molecular features detected only in "positive" features are automatically considered potentially phenotype-associated. For all molecular features detected in both “positive” and “negative” samples, a representative value over all “positive” and “negative” samples is calculated (the mean, median, or min-max scaled area or height; min-max where the lowest value is taken from "positive" samples and the highest from "negative" samples). The quotient is compared against a user-defined fold-change threshold, and only features surpassing this threshold are considered phenotype-associated. This approach helps retain features that may be linked to bioactivity but are present at sub-inhibitory concentrations in "negative" samples (i.e., those without a biological readout). Of note, this mode does not perform predictive modeling but instead filters out features unlikely to be associated with the observed phenotype.

##### Quantitative Data – Percentage-Based Measurements

FERMO accepts quantitative phenotype data following a “higher-is-better” logic, which is often the case for percentage-like data. Typically this data is generated at one specific sample concentration (e.g., measured bioactivity). FERMO supports multiple measurements per sample and allows multiple independent assays at different concentrations. The processing algorithm follows these steps:

1. Values below zero are set to zero.
2. Duplicate measurements per sample are averaged using either the mean or median.
3. The areas of molecular features detected in more than three samples are z-transformed along with the phenotype measurements.
4. Transformed feature areas and percentage values are correlated using Pearson correlation.
5. The resulting p-values are corrected for multiple hypothesis testing using a user-specified correction method (e.g. Bonferroni).
6. Features that exceed user-defined thresholds for both correlation coefficient and adjusted p-value are considered phenotype-associated.

##### Quantitative Data – Concentration-Based Measurements

FERMO also accepts quantitative phenotype data following a “lower-is-better” logic, which is often the case for concentration-based measurements (e.g. minimum inhibitory concentration (MIC) per sample). FERMO supports multiple measurements per sample and allows multiple independent assays at different concentrations. Of note, inactive samples must be assigned a value of 0, to distinguish them from the minimum concentration at which activity was still observed. The processing algorithm follows these steps:

1. Duplicate measurements per sample are averaged using either the mean or median.
2. The areas of molecular features detected in more than three samples are z-transformed.
3. MIC values are converted to their reciprocals (1 / measurement) or left as zero if the concentration was zero, then z-transformed.
4. Transformed feature areas and MIC values are correlated using Pearson correlation.

5. The resulting p-values are corrected for multiple hypothesis testing using a user-specified correction method (e.g. Bonferroni).
6. Features that exceed user-defined thresholds for both correlation coefficient and adjusted p-value are classified as bioactivity-associated.

#### Scores calculation

The calculation of feature- and sample-related scores summarizes the output of multiple modules in an easily filterable and searchable manner. Below is a description of the different scores utilized in FERMO.

##### Feature Phenotype Score

The Phenotype score summarizes phenotype information assignment, with higher scores indicating higher likeliness for the feature to be associated with the phenotype. The score is determined by the highest score determined by phenotype assignment if such an assignment was performed; else, it assigns zero. For more information on phenotype assignment, see the respective Methods section.

##### Feature Novelty Score

The Novelty score summarizes multiple annotations of feature identity, with higher scores indicating higher likeliness for the feature to be novel. If any annotation was performed, the highest matching score is subtracted from one; else, it assigns zero. For more information on feature annotation, see the respective Methods section.

##### Sample Mean Novelty Score

The mean of Novelty Scores of all molecular features detected in a sample.

##### Sample Diversity Score

The Diversity Score (*DS*) quantifies the chemical diversity observed in each sample relative to all other samples in the dataset. Spectral networks are used as a proxy for chemical families. For each spectral similarity networking algorithm (*i*), the *DS* is calculated as the ratio of the number of detected networks in the sample (*Ni*) to the total number of networks detected across all samples (*Ti*) for that algorithm (*Ni/Ti*). The highest ratio among all tested networking algorithms is retained as the final *DS*. A higher score indicates greater chemical uniqueness of the sample. Since diversity scores are influenced by networking parameters, they are only comparable within a single FERMO analysis run.

$$DS = max\,(Ni \,/\, Ti)$$

##### Sample Specificity Score

The Sample Specificity Score (*SS*) quantifies the proportion of chemical families unique to a given sample compared to all other samples. Again, similarity networks are used as a proxy for chemical families. For each spectral similarity networking algorithm (*i*), the $SS$ is calculated by subtracting sample-specific networks (*Si)* from networks detected across multiple samples (*Ni)*, and dividing by the total number of networks. The highest ratio among all tested networking algorithms is retained as the final $SS$. A higher score indicates a sample with more unique chemistry compared to other samples in the dataset. As specificity scores are influenced by networking parameters, they are only comparable within a single FERMO analysis run.

$$SS = max\left( (Ni - Si) / Ti \right)$$

#### Pseudo-chromatogram drawing

FERMO approximates extracted ion chromatograms (EICs) as pseudo-chromatograms. Unlike traditional EICs, which are continuous time–intensity traces, FERMO constructs pseudo-chromatograms from discrete data points extracted from the input peak table. These include the retention times at the start, apex, and end of each feature, as well as the full width at half maximum (FWHM). FERMO performs a series of sanity checks to identify and correct implausible values resulting from imperfect peak picking (e.g., an FWHM exceeding the duration between feature start and end), then generates a reconstructed retention time–intensity trace. This approach often yields pseudo-chromatograms that closely resemble the original data (see Supplementary Figure S1), though features with significant tailing or fronting may be less accurately represented. Early versions of the algorithm incorporated tailing and asymmetry factors, but these were later omitted due to their inconsistent availability in peak tables. Importantly, pseudo-chromatograms produced by FERMO are approximations intended primarily for visualization, and users are encouraged to consult the original data for confirmation.

### Supplementary Data

#### Supplementary Table S1: Collection of hyperlinks referenced in main text

| Column ID | Corresponding section in main manuscript | Description | Hyperlink | Zenodo |
|---|---|---|---|---|
| A | Hypothesis-driven prioritization using the FERMO dashboard | Feedback gathered during FERMO beta testing study in 2023 | https://github.com/fermo-metabolomics/fermo_ms/blob/main/supp_data/fermo_beta_testing_responses.csv | https://doi.org/10.5281/zenodo.15203137 |
| B | Hypothesis-driven prioritization using the FERMO dashboard | Online documentation of FERMO | https://fermo-metabolomics.github.io/fermo_docs/ | https://doi.org/10.5281/zenodo.15149101 |
| C | Benchmarking on Euphorbia dendroides dataset | Overview of predictions shared by Bioactivity-based molecular networking and FERMO in benchmarking on Euphorbia dendroides dataset | https://github.com/fermo-metabolomics/fermo_ms/blob/main/case_study_1/overlap_prediction_fermo_biomn.csv | https://doi.org/10.5281/zenodo.15203137 |
| D | Benchmarking on Euphorbia dendroides dataset | Comparison on effect of sample size on significance value between FERMO and BioMN | https://github.com/fermo-metabolomics/fermo_ms/blob/main/case_study_1/pearson_phenotype.ipynb | https://doi.org/10.5281/zenodo.15203137 |
| E | Benchmarking on Euphorbia dendroides dataset | FERMO-generated session file containing results. Used to create Figure 3. | https://github.com/fermo-metabolomics/fermo_ms/blob/main/case_study_1/fermo_analysis/results/out.fermo.session.json | https://doi.org/10.5281/zenodo.15203137 |
| F | Benchmarking FERMO against established phenotype-prioritization workflows | MZmine3 batch processing file | https://github.com/fermo-metabolomics/fermo_ms/blob/main/case_study_1/mzmine3_batch.xml | https://doi.org/10.5281/zenodo.15203137 |
| G | Benchmarking FERMO against established phenotype-prioritization workflows | FERMO processing parameters and results | https://github.com/fermo-metabolomics/fermo_ms/blob/main/case_study_1/fermo_analysis | https://doi.org/10.5281/zenodo.15203137 |
| H | FERMO effectively prioritizes bioactive actinomycin D in an OSMAC study | FERMO processing parameters and results | https://github.com/fermo-metabolomics/fermo_ms/blob/main/case_study_2/ | https://doi.org/10.5281/zenodo.15203137 |

#### Supplementary Table S2: Benchmarking of FERMO run time

| Nr features | Nr samples | Minimal processing | Data organization modules | Annotation modules (incl. score calc) | Time (sec) | Can be visualized in FERMO GUI |
|---|---|---|---|---|---|---|
| 619 | 82 | Y | Y | Y | 126 | Y |
| 3610 | 82 | Y | N | N | 36 | Y |
| 3610 | 82 | Y | Y | N | 248 | Y |
| 3610 | 82 | Y | Y | Y | 418 | Y |

*Subset of MassIVE (MSV000085376), pre-processed with Mzmine 4.5.37, processed using fermo_core version 0.6.2 on a machine running Ubuntu 20.04.6 LTS with 512GiB System memory Intel(R), with Xeon(R) Gold 6342 CPU @ 2.80GHz. Visualization was performed with FERMO GUI 1.0.9 in offline mode using Firefox v137.0.0.*

Supplementary Table S3: Qualitative comparison of metabolomics prioritization tools with FERMO

| Name | MS data format | Ionization | Orthogonal data | MS/MS organization | Annotation | Score calculation | Parameter management | Type application | Reference |
|---|---|---|---|---|---|---|---|---|---|
| FERMO | peak table, MS/MS data | +/- | phenotype, grouping, genomics | ✓ | ✓ | ✓ | ✓ | Web GUI/ CLI | This publication |
| Bioactivity-Based Molecular Networking | Peak table | +/- | phenotype | ✕ | ✕ | ✓ | ✕ | Jupyter notebook | Nothias et al 2018 |
| NP Analyst | peak table, raw data | +/- | phenotype | ✓ | ✓ | ✓ | ✕ | Web GUI | Lee et al 2022 |
| NP3 MS Workflow | raw data | + | phenotype, grouping | ✓ | ✓ | ✓ | ✕ | CLI | Bazzano et al 2022 |
| MetEx | peak table | +/- | elicitor library | ✕ | ✓ | ✓ | ✕ | Web GUI | Covington and Seyedsayamdost 2021 |
| GNPS | peak table, MS/MS data, raw data | +/- | grouping | ✓ | ✓ | ✕ | ✕ | Web GUI | Wang et al 2016 |
| Metabo Analyst | peak table, raw data | +/- | grouping, transcriptomic, proteomic | ✓ | ✓ | ✕ | ✕ | Web GUI/ CLI | Xia et al 2009 |

### Supplementary Table S4: FAIR4RS Checklist

| FAIR4RS principles[14] | Adherence | Note |
|---|---|---|
| F1. Software is assigned a globally unique and persistent identifier. | Y | Registered at bio.tools under biotools:fermo and biotools:fermo-core |
| F1.1. Components of the software representing levels of granularity are assigned distinct identifiers. | Y | FERMO (GUI) and fermo_core are managed distinctly |
| F1.2. Different versions of the software are assigned distinct identifiers. | Y | Version-controlled following Semantic Versioning |
| F2. Software is described with rich metadata. | Y | Registered on bio.tools |
| F3. Metadata clearly and explicitly include the identifier of the software they describe. | Y | Registered on bio.tools |
| F4. Metadata are FAIR, searchable and indexable. | Y | Registered on bio.tools |
| A1. Software is retrievable by its identifier using a standardised communications protocol. | Y | FERMO: https/browser, git<br>Fermo-core: https/browser, git, pip |
| A1.1. The protocol is open, free, and universally implementable. | Y | |
| A1.2. The protocol allows for an authentication and authorization procedure, where necessary. | N/A | |
| A2. Metadata are accessible, even when the software is no longer available. | Y | Registered on bio.tools |
| I1. Software reads, writes and exchanges data in a way that meets domain-relevant community standards. | Y | FERMO parses and produces Mzmine-generated peaktables (csv) |
| I2. Software includes qualified references to other objects. | Y | Crosslinks to PubChem/GNPS/MIBiG where applicable |
| R1. Software is described with a plurality of accurate and relevant attributes. | Y | Metadata in pyproject.toml files, metadata provided on bio.tools |
| R1.1. Software is given a clear and accessible license. | Y | MIT License |
| R1.2. Software is associated with detailed provenance. | Y | Version history on GitHub page |
| R2. Software includes qualified references to other software. | Y | Interoperable with existing tools |
| R3. Software meets domain-relevant community standards. | Y | Reads mgf files |

#### Supplementary Figure S1: Pseudo-chromatograms

Comparison of pseudo-chromatograms generated by FERMO with original total ion chromatograms as visualized by MZmine3.

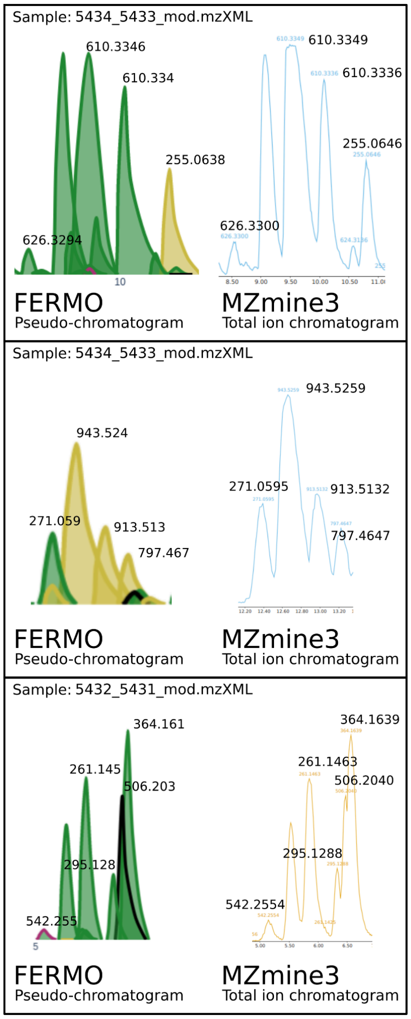


Original .mzXML files are available at https://github.com/fermo-metabolomics/fermo_ms/blob/main/supp_data

### References

1. Beck, K. *Test Driven Development: By Example*. (Addison-Wesley Professional, 2022).

2. Wang, M. *et al.* Sharing and community curation of mass spectrometry data with Global Natural Products Social Molecular Networking. *Nat Biotechnol* **34**, 828–837 (2016).

3. Watrous, J. *et al.* Mass spectral molecular networking of living microbial colonies. *Proc Natl Acad Sci U S A* **109**, E1743–52 (2012).

4. Huber, F., van der Burg, S., van der Hooft, J. J. J. & Ridder, L. MS2DeepScore: a novel deep learning similarity measure to compare tandem mass spectra. *J Cheminform* **13**, 84 (2021).

5. Kersten, R. D. *et al.* A mass spectrometry-guided genome mining approach for natural product peptidogenomics. *Nat Chem Biol* **7**, 794–802 (2011).

6. Kersten, R. D. *et al.* Glycogenomics as a mass spectrometry-guided genome-mining method for microbial glycosylated molecules. *Proc Natl Acad Sci U S A* **110**, E4407–16 (2013).

7. Niessen, W. M. A. & Correa C., R. A. *Interpretation of MS-MS Mass Spectra of Drugs and Pesticides*. (John Wiley & Sons, 2017).

8. de Jonge, N. F. *et al.* MS2Query: reliable and scalable MS mass spectra-based analogue search. *Nat Commun* **14**, 1752 (2023).

9. Blin, K. *et al.* antiSMASH 7.0: new and improved predictions for detection, regulation, chemical structures and visualisation. *Nucleic Acids Res* **51**, W46–W50 (2023).

10. Zdouc, M. M. *et al.* MIBiG 4.0: advancing biosynthetic gene cluster curation through global collaboration. *Nucleic Acids Res* **53**, D678–D690 (2025).

11. Strobel, M., Shahneh, M. R. Z., Gil de la Fuente, A., El Abiead, Y. & Wang, M. Tandem Mass Spectrometry Dataset for Machine Learning in Metabolomics. Zenodo https://doi.org/10.5281/ZENODO.11193898 (2024).

12. Terlouw, B. R. *et al.* MIBiG 3.0: a community-driven effort to annotate experimentally validated biosynthetic gene clusters. *Nucleic Acids Res* **51**, D603–D610 (2023).

13. Wang, F. *et al.* CFM-ID 4.0: More Accurate ESI-MS/MS Spectral Prediction and Compound Identification. *Anal Chem* **93**, 11692–11700 (2021).

14. Barker, M. *et al.* Introducing the FAIR Principles for research software. *Sci Data* **9**, 622 (2022).
